# Developmental and transcriptomic responses of Hawaiian bobtail squid early stages to ocean warming and acidification

**DOI:** 10.1101/2024.10.31.621237

**Authors:** E. Otjacques, J.R. Paula, E.G. Ruby, J.C. Xavier, M.J. McFall-Ngai, R. Rosa, C. Schunter

## Abstract

Cephalopods play a central ecological role across all oceans and realms. However, under the current climate crisis, their physiology and behaviour are impacted, and we are beginning to comprehend the effects of environmental stressors at a molecular level. Here, we study the Hawaiian bobtail squid (*Euprymna scolopes*), known for its specific binary symbiosis with the bioluminescent bacterium *Vibrio fischeri* acquired post-hatching. We aim to understand the response (i.e., developmental and molecular) of *E. scolopes* after the embryogenetic exposure to different conditions: i) standard conditions (control), ii) increased CO_2_ (ΔpH 0.4 units), iii) warming (+3ºC), or iv) a combination of the two treatments. We observed a decrease in hatching success across all treatments relative to the control. Using transcriptomics, we identified a potential trade-off in favour of metabolism and energy production, at the expense of development under increased CO_2_. In contrast, elevated temperature shortened the developmental time and, at a molecular level, showed signs of alternative splicing and the potential for RNA editing. The data also suggest that the initiation of the symbiosis may be negatively affected by these environmental drivers of change in the biosphere, although coping mechanisms by the animal may occur.

## 2 Introduction

Since the industrial revolution, oceans are becoming warmer, more acidic, and subject to extreme events such as marine heatwaves [1–4]. These changes in seawater conditions are known to impact marine organisms and communities [5–7], from physiology to behaviour [8– 14]. As the ocean changes and extreme events are expected to increase in strength and frequency due to the continuous increase of carbon dioxide (CO_2_) in the atmosphere [2,3,15,16], it is important to understand the biological response of species to such stressors.

Cephalopods play an important ecological role in marine ecosystems throughout all oceans and realms with a central position in trophic food webs [17–19]. They are also recognized as a keystone group for their economic importance in fisheries [17,20–22]. However, cephalopods are influenced by environmental changes [14], which can affect their physiology and behaviour, showing signs of reduced metabolic rates and activity levels [23], and impairment in predatory behaviour [24]. Moreover, deleterious effects can be observed at early developmental stages [14], disrupting cephalopod reproduction, embryonic development, and hatching success [25–27]. In fact, elevated CO_2_ levels showed a reduction in the number of eggs laid [28] and the mantle length [29] of squid hatchlings, and to increase the developmental time as well as reduce the hatching success in cephalopods [12,29]. Such responses in cephalopods were also observed with the combined exposure to increased CO_2_ and increased temperature [14,27,30].

To complement the current ecological and physiological knowledge on cephalopod species, a molecular approach is much needed since differential gene expression can be a major driver in phenotypic plasticity [31–33]. With the continuous pressure of climate change, various molecular responses are observed, where cephalopods present differential gene expression related, but not exclusively, to transcription factors and splicing activity after exposure to different temperatures [34]. Moreover, potential adaptation is also shown through the expression of ADAR (adenosine deaminase RNA specific) that is responsible for A-to-I RNA editing, with temperature playing a major role [34–36]. Finally, cephalopods exposed to higher CO_2_ concentrations present also molecular responses, linked to alterations in behaviour for example [37].

In this study, we investigate the transcriptomic response of the Hawaiian bobtail squid (*Euprymna scolopes*) exposed to increased CO_2_, elevated temperature and the combination of these two environmental factors, during embryonic development. *E. scolopes* is a small sepiolid species from the Hawaiian archipelago’s coastal waters, known for its binary symbiosis with the bioluminescent bacterium *Vibrio fischeri* [38]. Bobtail squids hatch without the symbiont and acquire the bacterial partner in the first hours post-hatching [39]. Whereas we have extensive knowledge of the animal’s relationship with the bacterial symbiont under standard laboratory conditions, environmental stress, such as seawater temperature or pH, has only been tested to understand the adaptation of *V. fischeri* in light of this symbiosis [40,41]. In contrast, the influences of environmental change on the bobtail squid host itself are poorly understood.

Here, we aim to understand the biological response of the Hawaiian bobtail squid *Euprymna scolopes*, after being exposed to different environmental conditions (i.e., increased CO_2_, warming and a combination of the two) during embryogenesis. Based on our knowledge of other cephalopods, we expect this species to present a lower hatching success across all treatment and reduced developmental time when exposed to warmer waters. By evaluating the transcriptomic response of this species, we aim to reveal the underlying molecular mechanisms of the response related to each treatment, expecting changes in developmental functions and metabolism. Understanding these changes in gene expression and the underlying functions allows the evaluation of the state of the bobtail squid early stages when exposed to near-future environmental changes.

## 3 Material and Methods

### 3.1 Experimental setup

In January 2022, adult Hawaiian bobtail squids were collected from Paikō peninsula (Oahu, USA) and maintained, as a breeding stock, in a flow-through system at the facilities of Kewalo Marine Laboratory (Oahu, USA). At the end of 4 months, a single clutch was prepared, packed in a temperature-insulated box, and shipped one day after being laid to the aquatic facility Laboratório Marítimo da Guia (Cascais, Portugal). The eggs were carefully separated and randomly distributed into 9 L plastic tanks (12 tanks in total, 3 replicates per treatment). These tanks were placed into two recirculating aquaria systems of approximately 92 L each, both separated into two water baths (4 WB in total, each corresponding to one treatment). As a semi-open system, the water in each WB was renewed by the constant addition of new water through a dripping system. After an acclimation of two days at control conditions, the eggs were reared until hatching in one of the following treatments: i) ‘control’ (25ºC; *p*CO_2_ = 320µatm, pH = 8.1), ii) ‘increased CO_2_’ (25ºC; *p*CO_2_ = 910 µatm, pH = 7.7), iii) ‘warming’ (28ºC; *p*CO_2_ = 320 µatm, pH = 8.1), and iv) ‘increased CO_2_ and warming’ (28ºC; pCO_2_ = 910 µatm, pH = 7.7). The ‘control’ temperature was based on the average water temperature observed in March and April in Oahu (i.e., 25ºC). Furthermore, the temperature and high CO_2_ were based on the IPCC’s RCP scenario 8.5 (i.e., +3 ºC; ΔpH = 0.4 units). Following the acclimation period, the water parameters were gradually altered to reach the final values for each treatment. Temperature was increased by +1 ºC per day and the pH lowered to approximately 0.1 unit per day through the injection of CO_2_ into the water.

Seawater was pumped directly from the ocean, filtered through a 1-µm mesh, and UV-sterilised (Vecton 120 Nano, TMC-Iberia, Lisbon, Portugal) before entering the aquatic systems. Filtration and UV-sterilisation systems in the experimental tanks and the control of seawater temperature and pH were performed following the methods described in Court et al., 2022. The photoperiod was kept under a 12h-light:12h-dark cycle using 8W LED lights.

Seawater parameters (Supplementary Table 1) were monitored daily using an oximeter VWR DO220 for oxygen levels and temperature (accuracy ± 1.5% and ± 0.3ºC, respectively), pH meter VWR pHenomenal for the pH (accuracy ± 0.005) and Hanna refractometer for the salinity (accuracy ± 1 PSU). The total alkalinity was measured weekly using a digital titrator (Sulfuric Acid 0.1600 N). The values of bicarbonate and *p*CO_2_ were subsequently calculated using the CO2SYS software.

### 3.2 Hatching success

To assess the hatching success at the end of the experiment, each egg capsule was examined under a scope to confirm the number of empty capsules (hatched individuals, n_total_hatched_ = 237) and the number of aborted embryos (n_total_aborted_ = 43) across treatments. Since the hatching success is represented by time-to-event data, we performed a survival analysis on this hatching success, according to the developmental time (i.e., number of days between eggs laid and hatching). More specifically, using R v. 4.3.3, the hatching success was assessed using the R package “survival” v. 3.6-4 [42], through a Cox proportional hazards regression model using the function “coxph”. The scaled residuals over time (Schoenfeld test; function “ggcoxzph”) were plotted to test the assumptions of the “coxph” model (proportional hazards, no over-influential observations and linearity of covariates). Since the requirements for the Schoenfeld test were not met, a non-parametric “survdiff” model was best fitted (Supplementary Figure 1, [43]). Moreover, post-hoc multiple comparisons were performed, and p-values were adjusted through Bonferroni–Hochberg corrections to avoid type I errors (Supplementary Table 2.A-B). Kaplan-Meier plots were created to illustrate the survival curves using the function “ggsurvplot” (R package “survminer” v. 0.4.9, [44]).

### 3.3 RNA extraction and RNA sequencing

Due to the lack of knowledge in the response of this species to climate change stressors and because hatchlings only measure around 2 mm, whole animals were used in the transcriptomic analysis. To understand the environmental response during the embryogenesis, animals were flash-frozen up to 2 h post-hatching and kept at -80ºC until RNA extractions. RNA was extracted using the AllPrep DNA/RNA Mini Kit (Qiagen), following the manufacturer’s protocol. Because hatching is usually triggered by a light cue [39], only the RNA of animals hatched 2 h after sunset (n = 8 per treatment) were tested for quality (Bioanalyzer) and further processed for sequencing by the Centre for PanorOmic Sciences of the University of Hong Kong. The sequencing libraries were prepared using the KAPA mRNA HyperPrep Kit, and Illumina NovaSeq 6000 was used for Pair-End 151 bp sequencing.

### 3.4 RNA-seq read processing

To understand the molecular basis after the embryogenesis exposure to the different treatments, an average of 66.6 million raw paired-end reads were processed using the following bioinformatic pipeline. The quality of reads after each processing step was inspected using FastQC v.0.11.9 [45]. The trimming of low quality reads and adapters was performed using

Trimmomatic v.0.39 [46] with the following parameters: ILLUMINACLIP:AllAdaptors.fa:2:30:15:8:true LEADING:3 TRAILING:3 SLIDINGWINDOW:4:20 MINLEN:40. To remove potential contamination, we used Kraken2 with a confidence of 0.5 [47], using the standard database from NCBI RefSeq as reference (version of the 05/06/2023), which contains libraries for archaea, bacteria, virus, plasmid, human and vectors [47]. Further filtration of low quality and short reads was performed using ‘filter_illumina’ script from DRAP [48]. Finally, reads from ribosomal RNA (rRNA) were identified and removed by performing a mapping of the sequences to the SILVA databases (SILVA_138_SSUParc_tax_silva.full_metadata.gz, SILVA_132_LSUParc.full_metadata.gz, [49], using bowtie2 v.2.4.1 [50] with very sensitive and local mode. The adapter-free, quality-trimmed, decontaminated and filtered paired-end reads (average 29.3 filtered pair-ended reads) were then mapped to the reference genome available for *Euprymna scolopes* [51] using STAR v.2.7.10b (parameters: --outSAMtype BAM Unsorted SortedByCoordinate --outFilterScoreMinOverLread 0.50 --outFilterMatchNminOverLread 0.50, [52]. On average, 77.03 ± 3.84% reads mapped to the reference genome (Supplementary Table 3). Raw read counts per gene were obtained using featureCounts v.2.0.6 [53]. Finally, a functional annotation was also performed using EggNOG-mapper v.2.1.10 [54].

### 3.5 Differential gene expression analysis

To understand the differential expression of genes between treatments, we used the R package DESeq2 v.1.40.2 [55] with a Wald test. We examined the count matrix for potential outliers. Therefore, after normalizing the variance of the count data, we performed a Principal Component Analysis (PCA), using a confidence level of 95%. Outliers were identified as samples outside the confidence ellipse of the PCA. Following this method, two samples were removed from the analysis (i.e., one from the ‘control’ treatment and one from the ‘warming’ treatment; Supplementary Figure 2). Moreover, low expression genes (< 10 read counts) were also excluded from the rest of the analysis. To obtain the list of differentially expressed genes (DEGs), we performed pairwise comparisons between each condition: i) ‘control’ vs. ‘increased CO_2_’, ii) control vs. ‘warming’, iii) ‘control’ vs. ‘increased CO_2_ and warming’. We identified DEGs with FDR adjusted p-value < 0.05 and a baseMean > 10. We used the log2Fold change as an additional criterion to decrease false positives considering significance only with absolute log2fold change > 0.3.

### 3.6 Weighted gene co-expression network analysis (WGCNA) and module eigengenes correlation to environmental traits

An additional analysis to study the correlation between gene expression and treatments was performed through the weighted gene co-expression network analysis. We normalized the count data and removed low read counts (< 10 counts in ≥ 90% samples) using DESeq2. Subsequently, we performed a step-by-step network construction and module detection using the WGCNA v. 1.72-5 R package [56]. The selection of the soft-threshold power (SFT) and the correlation network adjacency was calculated using 8 as the SFT (Supplementary figure 3).

The adjacency was transformed into a topological overlap matrix (TOM) and the corresponding dissimilarity was calculated (1-TOM). We produced a hierarchical clustering using the “average” method and, with the dissimilarity TOM, created a dendrogram containing the obtained cluster of genes. The modules were identified using a dynamic tree cut with the following parameters: minClusterSize = 100, deepSplit = 3 and pamRespectsDendro = FALSE. Modules with a similar expression profile were merged (branch height cut-off of 0.25 corresponding to a correlation of ≥ 0.75) and eigengenes were calculated for each module. These modules eigengenes (MEs) were correlated to each treatment (i.e., ‘warming’, ‘increased CO_2_’, and ‘increased CO_2_ and warming’) using the Pearson correlation test and a correlation heatmap was created (Supplementary figure 4). For a given correlation, student asymptomatic p-values were calculated displaying the correlation values of the modules for each trait. Only the significant modules displayed in the heatmap (p-value < 0.05) correlated to each trait were selected for further analysis.

### 3.7 Gene set enrichment analysis (GSEA)

The GSEA aims to understand if groups of genes that fulfil a similar function [gene ontology (GO)] showed significant and consistent differences between each treatment and the control conditions. We created an annotation data package specific for the Hawaiian bobtail squid *Euprymna scolopes* based on the GO terms for each gene described in the reference genome [51], using the R package AnnotationForge v. 1.42.2 [57]. Using the organism-specific annotation package created, we then performed a gene set enrichment analysis (GSEA) using the R package clusterProfiler v. 4.8.3 [58]. We performed the GSEA on the outputs from both the DESeq2 analysis and the significant modules from the WGCNA. Moreover, we performed additional GSEA on upregulated and downregulated genes under ‘increased CO_2_’ compared to ‘control’. No enrichment was found using the DEGs between ‘warming’ vs. ‘control’, nor the DEGs between ‘warming and increased CO_2_’ vs. ‘control’. Moreover, no enrichment was found in the modules *darkturquoise* (correlated to ‘increased CO_2_’), nor *grey* (correlated to ‘increased CO_2_ and warming’). All GSEA were performed using a minimum gene set size (GSS) of 10 and a maximum GSS of 500. Moreover, p-values were adjusted for multiple comparison using the method of “Benjamini-Hochberg” and a threshold of significant was set to padj < 0.05.

## 4 Results

### 4.1 Hatching success

We observed a significant decrease in hatching success in all treatments compared to control animals (Supplementary Table 2, 4). 96.4% of bobtail squids raised under control conditions hatched (n_hatching_control_/n_total_control_ = 54/56), however, there was a decrease to 78.5% in hatching success for the animals raised under increased CO_2_ conditions (n_hatching_increasedCO2_/n_total_increasedCO2_ = 62/79, p-value = 0.0023), to 85.9% under warming conditions (n_hatching_warming_/n_total_warming_ = 61/71, p-value < 0.001) and to 81.1% under the combination of increased CO_2_ and warming conditions (n_hatching_warming_/n_total_warming_ = 60/74, p-value = 0.0101; Figure 1). After comparing the hatching success between treatments, we observed that animals raised under increased CO_2_ conditions also exhibited lower hatching success compared to warming conditions (p-value < 0.001) and to the combination of increased CO_2_ and warming conditions (p-value < 0.001). However, animals raised in warming conditions did not have a significantly different hatching success compared to bobtail squids reared under the combination of increased CO_2_ and warming (p-value = 0.1891; Supplementary Table 2).

**Figure 1.**
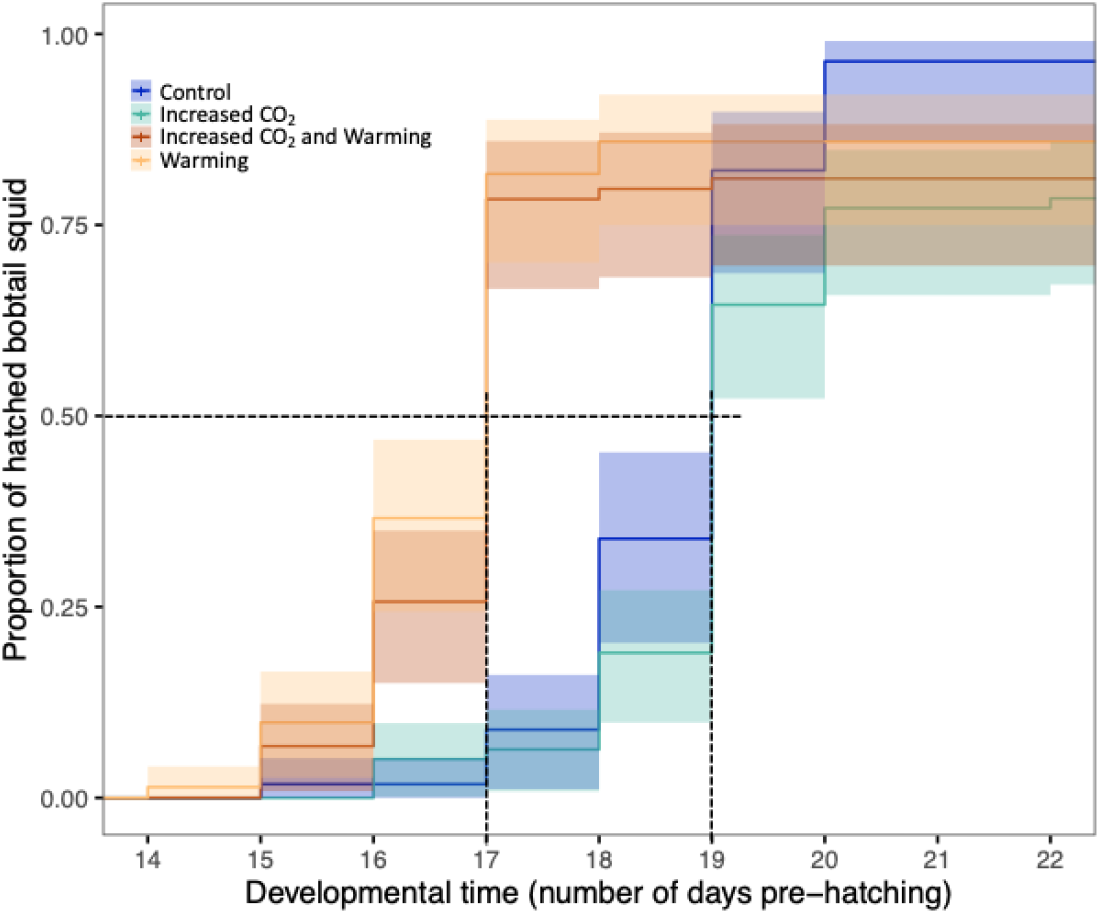
Hatching success of the Hawaiian bobtail squid reared under different environmental conditions. The proportion of hatched bobtail squid was measured according to the number of days pre-hatching (developmental time). The presence of hatching was verified daily. The Kaplan-Meier survival trajectories illustrate the survival trajectories according to each treatment (the colour code for each treatment is shown in the upper-left quadrant of the figure). The lines represent the rate of hatched bobtail squid, at each given day of exposure. The shaded area shows the 95% confidence intervals. The dashed lines show the hatching time, corresponding to the median developmental time.

Animals reared under control temperatures (i.e., ‘control’ and ‘increased CO_2_’) showed a hatching time (time when 50% of embryos hatched compared to an expected hatching of 100%, corresponds to the median developmental time) of 19 days. However, bobtail squids raised under warmer temperatures (i.e., ‘warming’, and ‘increased CO_2_ and warming’) hatched 2 d earlier (hatching time of 17 days; Figure 1, Supplementary Table 4).

### 4.2 Differentially expressed genes

By comparing the expression profile across the four treatments, we observed a higher variance in the ‘increased CO_2_’ treatment compared to the ‘control’ than with the other treatments (Figure 2). We identified a total of 970 differentially expressed genes (DEGs) between the ‘control’ and the ‘increased CO_2_’ treatments, 660 genes were upregulated and the remaining 310 were downregulated under the ‘increased CO_2_’ condition (Figure 3, Supplementary Table 5). On the other hand, a total of 21 DEGs were found between the ‘control’ and the ‘warming’ conditions, seven genes were upregulated, and 14 genes were downregulated with temperature (Figure 3, Supplementary Table 6). Finally, only three DEGs were found between the combined treatment (‘increased CO_2_ and warming’) and the ‘control’ condition; all three genes were downregulated under the combined treatment (Figure 3, Supplementary Table 7). Only one of the three DEGs was specific to the combined treatment and the other two downregulated genes were found under the ‘warming’ treatment alone (Figure 3).

**Figure 2.**
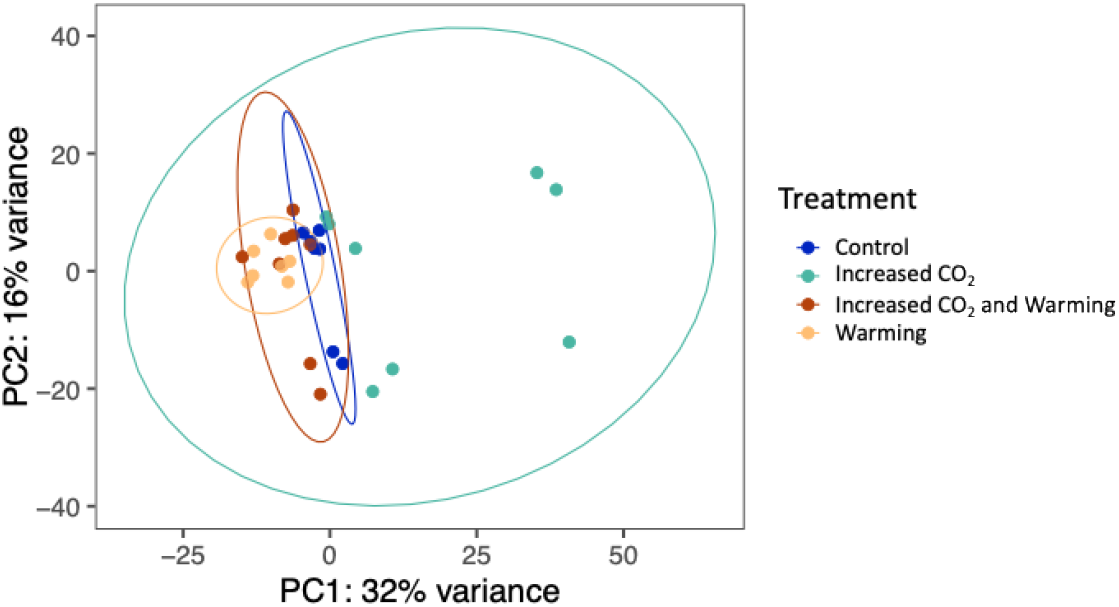
Principal component analysis on the normalized gene expression data. The ellipses represent the 95% confidence level, and the dots are the data points of each sample.

**Figure 3.**
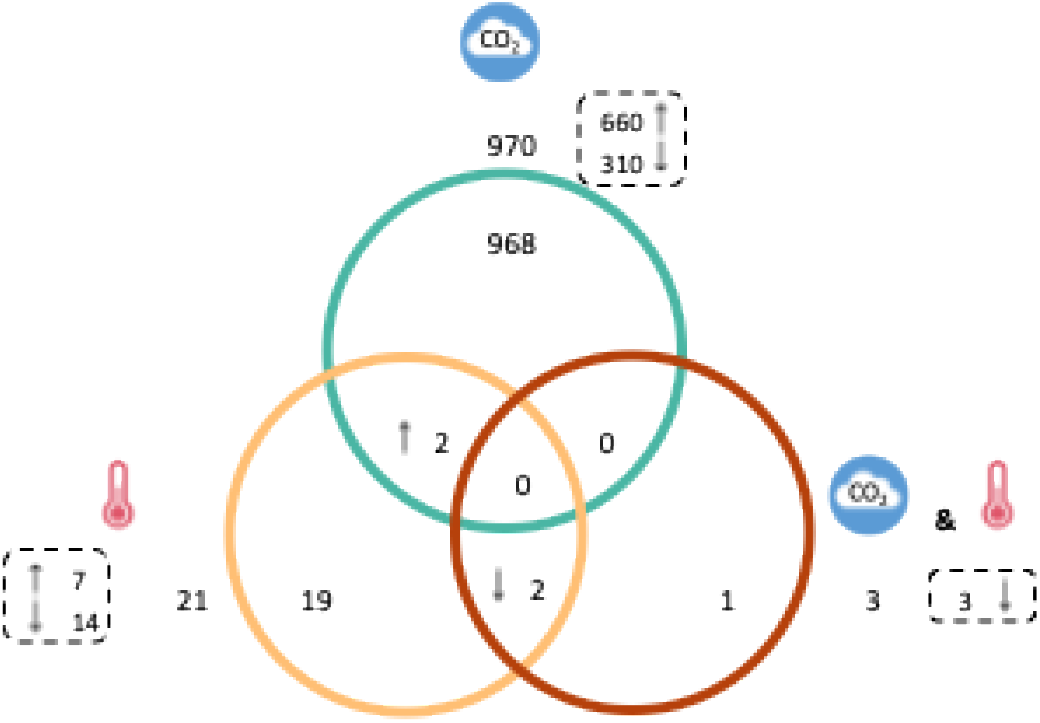
Venn diagram comparing the differentially expressed genes between the ‘control’ and each of the treatment. Green = ‘increased CO_2_’, yellow = ‘warming’, brown = ‘increased CO_2_ and warming’, = upregulated genes and = downregulated genes.

### 4.3 Transcriptomic response to increased CO_2_ exposure

Exposure to elevated CO_2_ provoked the largest number of differentially expressed genes compared to the other treatments (Figure 3, Supplementary Table 5). We identified seven key functions related to the DEGs and genes correlated to increased CO_2_: 1) protein folding and handling; 2) energy production and metabolism, including electron transport chain; 3) immune response; 4) vesicle organization and transportation, and neuronal development; 5) behaviour and neurotransmitters; 6) developmental processes, cell adhesion and structure organization; 7) signalling pathways (Supplementary Tables 8-12).

More specifically, protein folding involved DEGs such as heat shock proteins, prolyl isomerase (PPIase) and several prefoldin subunits as well as a DNA helicases “ATP dependent 5’ 3’ DNA helicase activity” (*ruvbl2*). Underlying the same function we also found genes specifically associated with endoplasmic reticulum (*erp29, emc3*). On the other hand, DEGs related to energy production and metabolism were identified as several subunits of the NADH:Ubiquinone Oxidoreductase complex (*nduf*). We also detected the differential expression of Cytochrome c oxidase subunits (*cox* genes), prohibitin, ATPase with H^+^ transport (*atp6*, also known as V-ATPase) and ATP synthase, involved in the electron transport chain, associated with ATP synthesis, and oxidative phosphorylation.

Gene upregulation under increased CO_2_ showed similar functions as described but we established immune response as an additional function. Such genes included the nicotinamide phosphoribosyltransferase (*naprt*) and 26s proteasome subunits (*psmd*). Moreover, the functions of neuronal development and vesicle organization and transportation (including “synaptic vesicle maturation” or “regulation of dendrite development”) were identified through the presence of positively correlated genes, including genes such as synaptoporin (*synpr*), syntaxin-binding proteins (*stx*) synaptosomal associated protein (*snap47*) or as a Rab GTPase activating protein (*rabgap1*). We also found a protocadherin (*pcdh15*) associated with the neuronal function of “visual perception”. Finally, the functions associated with neurotransmission and behaviour were also involved as seen through the positive correlation of receptors for dopamine (*drd2*), serotonin (*htr*), GABA_A_ and GABA_B_ (*gabra4* and *gabbr1*, respectively), and for glutamate (*grin2b*).

Downregulated and negatively correlated genes under increased CO_2_, on the other hand, were related to developmental processes (e.g., “endoderm and mesoderm formation and differentiation”, “gastrulation”, “ossification”, “striated muscle cell development”, etc.), but also genes for cell adhesion and structure (e.g., “cell-substrate adhesion”, “extracellular matrix organization”, etc.). Some of these genes were the transcriptional regulator β-catenin (*apc2*) involved in development, cadherin (*fat4*), or part of the *sox* family of transcription factors (e.g., *sox17*) involved in cell differentiation. We have also identified a negative correlation with genes coding for ryanodine (*ryr2*), fibroblast growth factor activated receptor (*fgfr4*) and genes for chains of collagen (*col*), involved in several aspect of muscle and embryonic development. Finally, the negatively correlated genes involved in signalling were more specifically belonging to the Wnt signalling pathway (*wnt*), which can be involved in developmental processes dependent on colonisation by microbes [59].

### 4.4 Transcriptomic response to increased temperature

Increased temperature did not induce a large response with only 21 DEGs (Figure 3, Supplementary Table 6). Two of these genes were also identified under increased CO_2_ and were upregulated: one could not be characterized, and the other was a gene coding for an Opioid growth factor receptor (*ogfr*). On the other hand, some temperature-specific DEGs were recognized as a calcium-activated potassium channel (*kcnn2*; upregulated with temperature), and a member of the molecular chaperone cytochrome p450 family (*cyp4v2*; downregulated).

In addition to the DEGs, we identified five main functions underlying gene networks correlated with temperature: 1) RNA processing and splicing; 2) metabolic and catabolic processes; 3) detoxification response; 4) reproductive processes; and 5) signalling pathways linked to the immune response (Supplementary Tables 13-14). RNA splicing was linked to positively correlated genes coding for splicing factors such as several serine/arginine rich splicing factors (*srsf*), and heterogenous nuclear ribonucleoprotein (*hnrnpu*). Other RNA processing functions were found through the expression of primary miRNA methylation (*mettl3*). We have also discovered a gene for an adenosine deaminase-like (*adal*), responsible for the adenosine catabolic process and inosine biosynthetic process. Positively correlated genes featuring metabolic and catabolic processes were characterized, but not only, as acetyltransferase and methyltransferase (*cat1* and *carnmt1*, involved in the “amino acid metabolic processes”) or as a sirtuin (*sirt4*).

On the other hand, we characterized other metabolic processes (for glutathione, fatty acid, prostaglandin and prostanoid) linked to negatively correlated genes. These genes included the glutathione-S-transferase (*gst*), and the thromboxane-A synthase 1 (*tbxas1*). Moreover, detoxification response (i.e., “response to reactive oxygen species”) also involved a negatively correlated gene related to the 3’,5’ cyclic GMP phosphodiesterase activity, which plays a role in the nitric oxide pathway. We also identified another gene for the glutathione-S-transferase (*hpgds*), recognized to be involved in reproductive processes such as “regulation of germ cell proliferation” or “male germ cell proliferation”. Finally, we found that other signalling pathways related to the immune response to be negatively correlated to temperature. We observed an enrichment in “TRIF-dependent toll-like receptor signalling pathway”, associated with the genes for the NF-κB essential modulator NEMO (*ikbkg*) and the protein tyrosine kinase *ikbke* also known as the “I-kappa-B kinase epsilon”. Both genes are involved in the NF-κB signalling pathway related to the immune response.

### 4.5 Transcriptomic response to the combined exposure of increased CO_2_ and increased temperature

Only three genes were differentially expressed under the combined treatment of ‘increased CO_2_ and warming’ (Figure 3, Supplementary Table 7). The oxidative stress induced growth inhibitor family member 2 (*osgin2*) was the only DEG specific to this treatment. The two other genes were also found to be significantly downregulated in the ‘warming’ treatment. They were identified as a “leucine-rich repeat-containing protein 74A-like” (*lrrc74a*), and the protein coding gene *ankar* (armadillo/β-catenin like repeats).

Functions that were positively correlated to ‘increased CO_2_ and warming’ (module *grey*; Supplementary Table 15), included immune response and signalling pathways with underlying genes such as a toll like receptor (*tlr2*, known for its role in the detection of microbes) and a member of the protein kinase family (*map3k7*, also involved in the NF-κB signalling pathway). Moreover, we found genes involved in RNA/DNA processing and repair as well as transcriptional regulation. We detected the protein coding gene *msl3* (i.e., a methylated histone binding), RNA polymerase I and II, reverse transcriptase, the *pif1* helicase responsible for DNA replication and repair, as well as the zinc finger transcription factor (*snai2*). Finally, we also found the influence of this treatment on neurotransmission, through the positive correlation of the glutamate receptor *gria2*.

## 5 Discussion

Although most cephalopods are known to be affected by climate change-related stressors, there is a profound lack of knowledge on the sepiolids response to environmental factors. Here, we show that increased temperature and CO_2_ are negatively impacting the hatching success of the Hawaiian bobtail squid, the latter exhibited the lowest hatching success of all treatments. This decreased hatching shows the vulnerability of this species to changes in pH, potentially due to the change in the acid-base balance during development and the function of ion regulatory structures [60]. Moreover, as a tropical species with less seasonal variation, we observed the Hawaiian bobtail squid is also sensitive to temperature with decreased hatching success, which is consistent with the decreased number of hatchling in other squid species [61], with some depending on the season [27]. Since our interpretation relies on a single clutch only, it could limit the variation in the data, but it may have also restricted the number of responses for this species. However, together with previous studies, it becomes clear that hatching and, therefore, the fitness of bobtail squids are likely impacted by near-future climate change.

Developmental time (i.e., the number of days pre-hatching) varies between cephalopod species and depends on the exposure to environmental stressors. Temperature always reveals itself as the main driver for a reduced developmental time, in contrast to increased CO_2_ exposure leading to an increase in developmental time [12,25,27,30,62,63]. Here, we show the divergence of bobtail squid compared to other cephalopods. Whereas bobtail squids developmental time showed the same reduction in time under elevated temperature, bobtail squid exposed to increased CO_2_ did not exhibit a longer developmental time. A shorter developmental time may be related to increased metabolic rates of embryos under elevated temperature, with an increased oxygen demand [25]. On the contrary, metabolic suppression is thought to explain a delayed hatching after the exposure to increased CO_2_ [29]. Therefore, while we show that bobtail squid may also increase their metabolism under warner temperature resulting in a shorter developmental time, we suggest that bobtail squid do not reduce their metabolism under increased CO_2_, leading to a similar developmental time as ‘control’.

Hatching success and time of development are direct, measurable and observable, responses of the animal. Molecular data and the transcriptional response can help us comprehend these responses, in addition to understand broader changes in the animal. Just as for hatching success ‘increased CO_2_’ provoked the largest molecular response of *E. scolopes*. Our transcriptomic data may indicate a trade-off in favour of metabolism and energy production, at the expense of development, which could explain the negative impacts on hatching success. Changes in seawater pH induce acid-base imbalances which can be compensated through ion regulation machineries by several species of fishes and cephalopod [60,64–66]. In fact, cephalopod can actively perform such regulation during embryogenesis [67]. We found that bobtail squids upregulate genes coding for several subunits of V-type H^+^-ATPases (VHA), which may be used in counteracting the impact of acid-base changes and are considered as a key machinery to cope with extracellular pH unbalance [68,69], including in early ontological stages of cephalopods [70]. Although the implication of VHA as a response of stress-induced acid-base unbalance should be further characterized, the upregulation of these genes shows their potential involvement in coping with ocean acidification (i.e., ‘increased CO_2_). However, the regulation of the acid-base balance requires the consumption of energy and a coordination with the metabolism [66,71,72]. Hence, in response to increased CO_2_, it is not surprising to find large upregulation of cytochrome-c-oxidase (*cox*) and NADH dehydrogenase in the bobtail squid, as in many invertebrates (e.g., oyster [73–75], sea snail [76], spider crab [77], mussel [78]). Moreover, we found upregulation of prohibitin (PHB) when exposed to ‘increased CO_2_’, similar to that reported in the Pacific oyster [74]. PHB is a highly conserved protein across organisms, including marine vertebrates and invertebrates, that can be associated with the mitochondria [79,80]. We show that, with exposure to ‘increased CO_2_’, there is an upregulation of genes involved in metabolism and energy production, potentially indicating an increased demand of energy needed for acid base regulation.

On the other hand, we observed a downregulation and negative correlation of genes involved in development and cellular structure in response to ‘increased CO_2_’. Whereas β-catenin play a central role in the Wnt signalling pathway and the cadherin complex [81], Wnts are signalling proteins implicated in animal development [82,83], and recognized as important for cellular differentiation and organization [84]. On the other hand, the cadherin complex provides structural integrity and cell-cell adhesion [84,85]. Here, we show a coordinated negative response in Wnt, β-catenin and cadherin, which is consistent to the general downregulation of such genes in the Pacific oyster, mussels and corals under ocean acidification [86–89]. A global downregulation and negative correlation of these three components (i.e., Wnt, cadherin and β-catenin) under ‘increased CO_2_’, accompanied by adverse impacts on the hatching success, suggest a negative impact of future CO_2_ levels on the embryonic development of bobtail squids.

Elevated temperature exhibited a positive response in catabolic processes, as we observed a positive correlation to sirtuin. Sirtuins are NAD^+^-dependent deacylases involved in cellular stress response, conserved amongst vertebrates and invertebrates [90,91]. The sirtuin 4 (*sirt4*), in particular, codes for a mitochondrial protein [90] involved in the regulation of reactive oxygen species (ROS) production [92] which can reduce mitochondria dysfunction in mammalian cells and releasing the stress induced by oxidative stress [92]. Our results may indicate a positive response against heat stress and is consistent with the response of other organisms showing the importance of sirtuins in the regulation of cellular stress response [91,93,94]. Another effect of increased temperatures was found through the positive correlation of genes involved in RNA processing and splicing involving the spliceosome. This may indicate potential for plasticity and adaptation under heat stress [95]. Alternative splicing (AS) is deemed important for gene regulation, playing a role in tissue development and involve proteins acting in opposite ways [96,97]. We show a positive correlation of protein-coding genes for serine/arginine splicing factors (i.e., *srsf*), referenced as “splicing activators” and responsible in exon recognition [96], accompanied by the heterogenous ribonucleoprotein (i.e., *hnrnpu*), a “splicing repressor” which blocks the access of the spliceosome [96]. Through the positive expression of both activator and repressor of AS, we suggest bobtail squids to be capable of fine adjustments in AS with temperature, wherein an increase in AS is found as a response after stress exposure, like corals after exposure to marine heatwaves [95] or shrimps under high alkalinity [98]. Moreover, we also found differential expression in an adenosine deaminase-like gene (*adal*). Adenosine deaminase is an enzyme responsible for the RNA editing of Adenosine-to-Inosine (A-to-I) and is recognized as the most common RNA modification [99,100]. RNA editing events are known to be abundant in cephalopods [101]. In fact, it was found that RNA editing in an octopus was temperature dependent, in this case there was an increase in RNA editing with colder temperature [36]. Although an increased in temperature did not elicit major changes in gene expression per se, we show that it led to molecular responses that included the regulation of ROS from the positive correlation with sirtuins. Moreover, while further investigation into the extent of splicing patterns and RNA edited sites is needed, the positive correlation of *srsf, hnrnpu* and *adal* to increasing temperature in bobtail squid may indicate a potential for diversifying mRNA through AS and RNA editing in this species, which could lead to phenotypic plasticity.

In contrast, we identified genes related to the immune response to be negatively correlated with elevated temperature. More specifically, we show the negative correlation of protein coding genes *ikbkg* (coding for IKKγ/NEMO) and *map3k7* (coding for the protein also known as TAK1), which are both implicated in the activation of the NF-κB pathway [102– 104]. It is suggested that *E. scolopes* uses critical components of the NF-κB pathway (i.e., IKKγ) during the initiation of the symbiosis with the bacterial symbiont *Vibrio fischeri* [105]. Because of the negative correlation of the expression of such genes (i.e., ikbkg and *map3k7*) with temperature, we hypothesize that the colonisation of the bobtail squid, and subsequently the initiation of the symbiosis, may be negatively affected by increased temperature. Although the NF-κB pathway was negatively correlated with increasing temperature, this was not the case with exposure to the combination of treatments, since the *map3k7* was positively correlated to increased CO_2_ and warming combined. This finding is consistent with the enrichment of the MAPK signalling pathway in a cuttlefish, when exposed to combined high temperature and low pH [106]. Under the same combined treatment, an additional protein was found through the expression of *tlr2*, a Toll-like receptor, also implicated in the microbial detection and the Toll/NF-κB pathways [105]. Therefore, increased CO_2_ and temperature might have antagonist effects in relation to the immune response. Although future investigations in understanding the colonisation efficiency of hatchlings when exposed to these stressors is needed, we show that temperature may negatively affect the initiation of the symbiosis, but not the combined treatment.

In summary, we show how environmental stressors induced a general adverse biological response in the Hawaiian bobtail squid, with a decrease in hatching success overall. We indicate that temperature was the main driver of the reduced developmental time, while increased CO_2_ exhibited the strongest molecular response. We identify a trade-off between metabolism and energy production against development when exposed to increased CO_2_, which may explain the lowest hatching success in this treatment. Increased temperature induced a heat stress response implicating the regulation of ROS and RNA processing. In fact, as a response to temperature, bobtail squid may alter their RNA through alternate splicing and RNA editing, which may lead to phenotypic plasticity. Finally, we show that the symbiosis initiation between the bobtail squid and its bioluminescent symbiont may be altered with increasing temperatures, but not when exposed to combined increased CO_2_ and temperature. Daily variation in coastal seawater temperature may explain the different responses towards plasticity and variability under increased temperature [107,108]. Such responses may also apply to other coastal cephalopod species including sepiolids; environmental changes could for example alter the colonisation of *Sepiola spp*., which implicates two bacterial symbionts that have different temperature growth optimum [109]. While future investigations should include testing for RNA editing and influence on animal-bacteria symbiosis, our results show that development is affected in early life stages of bobtail squids, whereas there are also signs of increased phenotypic plasticity in response to environmental stressors.

## 6 Ethics

This research was conducted in compliance with the Portuguese and EU legislations on the protection of animals used for scientific purposes (Decreto-Lei 113/2013 and Directive 2010/63/EU, respectively).

## 7 Data accessibility

All datasets and R codes will be made publicly available in the Figshare repository upon publication.

The raw sequencing data will be made publicly available in NCBI upon publication.

## 8 Declaration of AI use

We have not used AI-assisted technologies in creating this article.

## 9 Authors’ contribution

Conceptualization: EO, CS; Data curation: EO; Formal analysis: EO; Funding acquisition: EGR, MMN, RR, CS; Investigation: EO; Methodology: EO, JRP; Project administration: CS; Resources: EGR, MMN, RR, CS; Supervision: JCX, MMN, RR, CS; Visualization: EO; Writing – original draft: EO, CS; Writing – review & editing: EO, JRP, EGR, JCX, MMN, RR, CS.

All authors gave final approval for the submission of this manuscript

## 12 Acknowledgments

We thank all members of the Laboratório Marítimo da Guia and Rui Rosa Lab, Schunter lab and McFall-Ngai–Ruby labs for their assistance in maintaining the aquatic system, collecting the animals at hatching and advice on data processing.

## References

[1] Hobday AJ, Oliver ECJ, Sen Gupta A, Benthuysen JA, Burrows MT, Donat MG, et al. Categorizing and naming marine heatwaves. Oceanography 2018;31:162–73.

[2] Bindoff NL, Cheung WWL, Kairo JG, Arístegui J, Guinder VA, Hallberg R, et al. Changing Ocean, Marine Ecosystems, and Dependent Communities. In: Pörtner HO, Roberts DC, Masson-Delmotte V, Zhai P, Tignor M, Poloczanska E, et al., editors. IPCC Spec. Rep. Ocean Cryosphere Chang. Clim., 2019, p. 142.

[3] Fox-Kemper B, Hewitt HT, Xiao C, Aðalgeirsdóttir G, Drijfhout SS, Edwards TL, et al. Ocean, Cryosphere and Sea Level Change. In: Masson-Delmotte V, Zhai P, Pirani A, Connors SL, Péan C, Berger C, et al., editors. Clim. Change 2021 Phys. Sci. Basis Contrib. Work. Group Sixth Assess. Rep. Intergov. Panel Clim. Change, Cambridge University Press; 2021, p. 152.

[4] Calvin K, Dasgupta D, Krinner G, Mukherji A, Thorne PW, Trisos C, et al. IPCC, 2023: Climate Change 2023: Synthesis Report. Contribution of Working Groups I, II and III to the Sixth Assessment Report of the Intergovernmental Panel on Climate Change [Core Writing Team, H. Lee and J. Romero (eds.)]. IPCC, Geneva, Switzerland. First. Intergovernmental Panel on Climate Change (IPCC); 2023. 10.59327/IPCC/AR6-9789291691647.

[5] Shirayama Y, Thornton H. Effect of increased atmospheric CO2 on shallow water marine benthos. J Geophys Res 2005;110:1–7. 10.1029/2004JC002618.

[6] Hoegh-Guldberg O, Mumby PJ, Hooten AJ, Steneck RS, Greenfield P, Gomez E, et al. Coral reefs under rapid climate change and ocean acidification. Science 2007;318:1737– 42. 10.1126/science.1152509.

[7] Barnes D, Peck L. Vulnerability of Antarctic shelf biodiversity to predicted regional warming. Clim Res 2008;37:149–63. 10.3354/cr00760.

[8] Wittmann AC, Pörtner HO. Sensitivities of extant animal taxa to ocean acidification. Nat Clim Change 2013;3:995–1001. 10.1038/nclimate1982.

[9] Repolho T, Duarte B, Dionísio G, Paula JR, Lopes AR, Rosa IC, et al. Seagrass ecophysiological performance under ocean warming and acidification. Sci Rep 2017;7:1–12. 10.1038/srep41443.

[10] Cattano C, Claudet J, Domenici P, Milazzo M. Living in a high CO2 world: a global meta-analysis shows multiple trait-mediated fish responses to ocean acidification. Ecol Monogr 2018;88:320–35. 10.1002/ecm.1297.

[11] Paula JR, Repolho T, Pegado MR, Thörnqvist PO, Bispo R, Winberg S, et al. Neurobiological and behavioural responses of cleaning mutualisms to ocean warming and acidification. Sci Rep 2019;9:1–10. 10.1038/s41598-019-49086-0.

[12] Otjacques E, Repolho T, Paula JR, Simão S, Baptista M, Rosa R. Cuttlefish buoyancy control in response to food availability and ocean acidification. Biology 2020;9. 10.3390/biology9070147.

[13] Shodipo MO, Duong B, Graba-Landry A, Grutter AS, Sikkel PC. Effect of acute seawater temperature increase on the survival of a fish ectoparasite. Oceans 2020;1:215–36. 10.3390/oceans1040016.

[14] Borges FO, Sampaio E, Santos CP, Rosa R. Climate-change impacts on cephalopods: A meta-analysis. Integr Comp Biol 2023;63:1240–65. 10.1093/icb/icad102.

[15] Frölicher TL. Extreme climatic events in the ocean. In: Cisneros-Montemayor A, Cheung WWL, Ota Y, editors. Predict. Future Oceans Sustain. Ocean Hum. Syst. Glob. Environ. Change, 2019, p. 53–60.

[16] Oliver ECJ, Benthuysen JA, Darmaraki S, Donat MG, Hobday AJ, Holbrook NJ, et al. Marine heatwaves. Annu Rev Mar Sci 2021;13:313–42. 10.1146/annurev-marine-032720-095144.

[17] Clarke MR. The role of cephalopods in the world’s oceans: general conclusions and the future. Philos Trans R Soc Lond B Biol Sci 1996;351:1105–12. 10.1098/rstb.1996.0096.

[18] de la Chesnais T, Fulton EA, Tracey SR, Pecl GT. The ecological role of cephalopods and their representation in ecosystem models. Rev Fish Biol Fish 2019;29:313–34. 10.1007/s11160-019-09554-2.

[19] Murphy KJ, Pecl GT, Richards SA, Semmens JM, Revill AT, Suthers IM, et al. Functional traits explain trophic allometries of cephalopods. J Anim Ecol 2020;89:2692–703. 10.1111/1365-2656.13333.

[20] Arkhipkin AI, Rodhouse PGK, Pierce GJ, Sauer W, Sakai M, Allcock L, et al. World squid fisheries. Rev Fish Sci Aquac 2015;23:92–252. 10.1080/23308249.2015.1026226.

[21] Xavier JC, Allcock AL, Cherel Y, Lipinski MR, Pierce GJ, Rodhouse PGK, et al. Future challenges in cephalopod research. J Mar Biol Assoc U K 2015;95:999–1015. 10.1017/S0025315414000782.

[22] Sauer WHH, Gleadall IG, Downey-Breedt N, Doubleday Z, Gillespie G, Haimovici M, et al. World octopus fisheries. Rev Fish Sci Aquac 2021;29:279–429. 10.1080/23308249.2019.1680603.

[23] Rosa R, Seibel BA. Synergistic effects of climate-related variables suggest future physiological impairment in a top oceanic predator. Proc Natl Acad Sci U S A 2008;105:20776–80.

[24] Spady BL, Munday PL, Watson SA. Predatory strategies and behaviours in cephalopods are altered by elevated CO2. Glob Change Biol 2018;24:2585–96. 10.1111/gcb.14098.

[25] Rosa R, Pimentel MS, Boavida-Portugal J, Teixeira T, Trübenbach K, Diniz M. Ocean warming enhances malformations, premature hatching, metabolic suppression and oxidative stress in the early life stages of a keystone squid. PLoS ONE 2012;7. 10.1371/journal.pone.0038282.

[26] Rosa R, Trubenbach K, Repolho T, Pimentel M, Faleiro F, Boavida-Portugal J, et al. Lower hypoxia thresholds of cuttlefish early life stages living in a warm acidified ocean. Proc R Soc B Biol Sci 2013;280:1–7. 10.1098/rspb.2013.1695.

[27] Rosa R, Trubenbach K, Pimentel MS, Boavida-Portugal J, Faleiro F, Baptista M, et al. Differential impacts of ocean acidification and warming on winter and summer progeny of a coastal squid (Loligo vulgaris). J Exp Biol 2014;217:518–25. 10.1242/jeb.096081.

[28] Spady BL, Munday PL, Watson S-A. Elevated seawater pCO2 affects reproduction and embryonic development in the pygmy squid, Idiosepius pygmaeus. Mar Environ Res 2019;153. 10.1016/j.marenvres.2019.104812.

[29] Kaplan MB, Mooney TA, McCorkle DC, Cohen AL. Adverse effects of ocean acidification on early development of squid (Doryteuthis pealeii). PLoS ONE 2013;8. 10.1371/journal.pone.0063714.

[30] Court M, Paula JR, Macau M, Otjacques E, Repolho T, Rosa R, et al. Camouflage and exploratory avoidance of newborn cuttlefish under warming and acidification. Biology 2022;11:1394. 10.3390/biology11101394.

[31] Schlichting CD, Smith H. Phenotypic plasticity: linking molecular mechanisms with evolutionary outcomes. Evol Ecol 2002;16:189–211. 10.1023/A:1019624425971.

[32] Logan ML, Cox CL. Genetic constraints, transcriptome plasticity, and the evolutionary response to climate change. Front Genet 2020;11:538226. 10.3389/fgene.2020.538226.

[33] Strader ME, Wong JM, Hofmann GE. Ocean acidification promotes broad transcriptomic responses in marine metazoans: a literature survey. Front Zool 2020;17:7. 10.1186/s12983-020-0350-9.

[34] Fuentes PP. Integrating physiology, behaviour and molecular mechanisms to understand impacts of ocean warming on southern calamari (Sepioteuthis australis). University of Tasmania, 2021.

[35] Garrett SC, Rosenthal JJC. A role for A-to-I RNA editing in temperature adaptation. Physiology 2012;27:362–9. 10.1152/physiol.00029.2012.

[36] Birk MA, Liscovitch-Brauer N, Dominguez MJ, McNeme S, Yue Y, Hoff JD, et al. Temperature-dependent RNA editing in octopus extensively recodes the neural proteome. Cell 2023;186:2544-2555.e13. 10.1016/j.cell.2023.05.004.

[37] Thomas JT, Huerlimann R, Schunter C, Watson S-A, Munday PL, Ravasi T. Transcriptomic responses in the nervous system and correlated behavioural changes of a cephalopod exposed to ocean acidification. BMC Genomics 2024;25. 10.1186/s12864-024-10542-5.

[38] Nyholm SV, McFall-Ngai MJ. A lasting symbiosis: how the Hawaiian bobtail squid finds and keeps its bioluminescent bacterial partner. Nat Rev Microbiol 2021;19:666–79. 10.1038/s41579-021-00567-y.

[39] Nyholm SV, McFall-Ngai MJ. The winnowing: Establishing the squid–Vibrio symbiosis. Nat Rev Microbiol 2004;2:632–42. 10.1038/nrmicro957.

[40] Cohen ML, Mashanova EV, Rosen NM, Soto W. Adaptation to temperature stress by Vibrio fischeri facilitates this microbe’s symbiosis with the Hawaiian bobtail squid (Euprymna scolopes). Evolution 2019;73:1885–97. 10.1111/evo.13819.

[41] Cohen ML, Mashanova EV, Jagannathan SV, Soto W. Adaptation to pH stress by Vibrio fischeri can affect its symbiosis with the Hawaiian bobtail squid (Euprymna scolopes). Microbiology 2020;166:262–77. 10.1099/mic.0.000884.

[42] Therneau T. A Package for Survival Analysis in R. R package 2024.

[43] Therneau TM, Grambsch PM. Modeling Survival Data: Extending the Cox Model. New York, NY: Springer New York; 2000. 10.1007/978-1-4757-3294-8.

[44] Kassambara A, Kosinski M, Biecek P. survminer: Drawing Survival Curves using “ggplot2”. R package. 2016:0.4.9.

[45] Andrews S. FastQC: A quality control tool for high throughput sequence data. https://www.bioinformatics.babraham.ac.uk/projects/fastqc/ 2010.

[46] Bolger AM, Lohse M, Usadel B. Trimmomatic: a flexible trimmer for Illumina sequence data. Bioinformatics 2014;30:2114–20. 10.1093/bioinformatics/btu170.

[47] Wood DE, Lu J, Langmead B. Improved metagenomic analysis with Kraken 2. Genome Biol 2019;20:257. 10.1186/s13059-019-1891-0.

[48] Cabau C, Escudié F, Djari A, Guiguen Y, Bobe J, Klopp C. Compacting and correcting Trinity and Oases RNA-Seq de novo assemblies. PeerJ 2017;5:e2988. 10.7717/peerj.2988.

[49] Quast C, Pruesse E, Yilmaz P, Gerken J, Schweer T, Yarza P, et al. The SILVA ribosomal RNA gene database project: improved data processing and web-based tools. Nucleic Acids Res 2012;41:D590–6. 10.1093/nar/gks1219.

[50] Langmead B, Salzberg SL. Fast gapped-read alignment with Bowtie 2. Nat Methods 2012;9:357–9. 10.1038/nmeth.1923.

[51] Rogers TF, Yalçın G, Briseno J, Vijayan N, Nyholm SV, Simakov O. Gene modelling and annotation for the Hawaiian bobtail squid, Euprymna scolopes. Sci Data 2024;11:40. 10.1038/s41597-023-02903-8.

[52] Dobin A, Davis CA, Schlesinger F, Drenkow J, Zaleski C, Jha S, et al. STAR: ultrafast universal RNA-seq aligner. Bioinformatics 2013;29:15–21. 10.1093/bioinformatics/bts635.

[53] Liao Y, Smyth GK, Shi W. featureCounts: an efficient general purpose program for assigning sequence reads to genomic features. Bioinformatics 2014;30:923–30. 10.1093/bioinformatics/btt656.

[54] Cantalapiedra CP, Hernández-Plaza A, Letunic I, Bork P, Huerta-Cepas J. eggNOG-mapper v2: Functional annotation, orthology assignments, and domain prediction at the metagenomic scale. Mol Biol Evol 2021;38:5825–9. 10.1093/molbev/msab293.

[55] Love MI, Huber W, Anders S. Moderated estimation of fold change and dispersion for RNA-seq data with DESeq2. Genome Biol 2014;15:550. 10.1186/s13059-014-0550-8.

[56] Langfelder P, Horvath S. WGCNA: an R package for weighted correlation network analysis. BMC Bioinformatics 2008;9:559. 10.1186/1471-2105-9-559.

[57] Carlson M, Pagès H. AnnotationForge: Tools for building SQLite-based annotation data packages. R package 2024.

[58] Yu G, Wang L-G, Han Y, He Q-Y. clusterProfiler: an R Package for Comparing Biological Themes Among Gene Clusters. OMICS J Integr Biol 2012;16:284–7. 10.1089/omi.2011.0118.

[59] McFall-Ngai M, Bosch TCG. Animal development in the microbial world: The power of experimental model systems. Curr. Top. Dev. Biol., vol. 141, Elsevier; 2021, p. 371–97. 10.1016/bs.ctdb.2020.10.002.

[60] Hu MY, Tseng YC, Stumpp M, Gutowska MA, Kiko R, Lucassen M, et al. Elevated seawater pCO2 differentially affects branchial acid-base transporters over the course of development in the cephalopod Sepia officinalis. Am J Physiol - Regul Integr Comp Physiol 2011;300:1100–14. 10.1152/ajpregu.00653.2010.

[61] Zakroff CJ, Mooney TA. Antagonistic interactions and clutch-dependent sensitivity induce variable responses to ocean acidification and warming in squid (Doryteuthis pealeii) embryos and paralarvae. Front Physiol 2020;11:501. 10.3389/fphys.2020.00501.

[62] Repolho T, Baptista M, Pimentel MS, Dionísio G, Trübenbach K, Lopes VM, et al. Developmental and physiological challenges of octopus (Octopus vulgaris) early life stages under ocean warming. J Comp Physiol [B] 2014;184:55–64. 10.1007/s00360-013-0783-y.

[63] Zakroff C, Mooney TA, Berumen ML. Dose-dependence and small-scale variability in responses to ocean acidification during squid, Doryteuthis pealeii, development. Mar Biol 2019;166:1–24. 10.1007/s00227-019-3510-8.

[64] Seidelin M, Brauner CJ, Jensen FB, Madsen SS. Vacuolar-Type H+ -ATPase and Na+, K+-ATPase Expression in Gills of Atlantic Salmon (Salmo salar) during Isolated and Combined Exposure to Hyperoxia and Hypercapnia in Fresh Water. Zoolog Sci 2001;18:1199–205. 10.2108/zsj.18.1199.

[65] Choe KP, Evans DH. Compensation for hypercapnia by a euryhaline elasmobranch: Effect of salinity and roles of gills and kidneys in fresh water. J Exp Zoolog A Comp Exp Biol 2003;297A:52–63. 10.1002/jez.a.10251.

[66] Gutowska MA, Melzner F, Langenbuch M, Bock C, Claireaux G, Pörtner HO. Acid-base regulatory ability of the cephalopod (Sepia officinalis) in response to environmental hypercapnia. J Comp Physiol [B] 2010;180:323–35. 10.1007/s00360-009-0412-y.

[67] Hu MY, Tseng Y-C, Lin L-Y, Chen P-Y, Charmantier-Daures M, Hwang P-P, et al. New insights into ion regulation of cephalopod molluscs: a role of epidermal ionocytes in acid-base regulation during embryogenesis. Am J Physiol-Regul Integr Comp Physiol 2011;301:R1700–9. 10.1152/ajpregu.00107.2011.

[68] Forgac M. Vacuolar ATPases: rotary proton pumps in physiology and pathophysiology. Nat Rev Mol Cell Biol 2007;8:917–29. 10.1038/nrm2272.

[69] Brown D, Paunescu TG, Breton S, Marshansky V. Regulation of the V-ATPase in kidney epithelial cells: dual role in acid–base homeostasis and vesicle trafficking. J Exp Biol 2009;212:1762–72. 10.1242/jeb.028803.

[70] Hu MY, Guh Y-J, Stumpp M, Lee J-R, Chen R-D, Sung P-H, et al. Branchial NH4+-dependent acid–base transport mechanisms and energy metabolism of squid (Sepioteuthis lessoniana) affected by seawater acidification 2014.

[71] Pörtner HO. Coordination of metabolism, acid-base regulation and haemocyanin function in cephalopods. Mar Freshw Behav Physiol 1995;25:131–48. 10.1080/10236249409378913.

[72] Dubyak GR. Ion homeostasis, channels, and transporters: an update on cellular mechanisms. Adv Physiol Educ 2004;28:143–54. 10.1152/advan.00046.2004.

[73] Thompson EL, O’Connor W, Parker L, Ross P, Raftos DA. Differential proteomic responses of selectively bred and wild-type Sydney rock oyster populations exposed to elevated CO2. Mol Ecol 2015;24:1248–62. 10.1111/mec.13111.

[74] Timmins-Schiffman E, Coffey WD, Hua W, Nunn BL, Dickinson GH, Roberts SB. Shotgun proteomics reveals physiological response to ocean acidification in Crassostrea gigas. BMC Genomics 2014;15.

[75] Wei L, Wang Q, Wu H, Ji C, Zhao J. Proteomic and metabolomic responses of Pacific oyster Crassostrea gigas to elevated pCO2 exposure. J Proteomics 2015;112:83–94. 10.1016/j.jprot.2014.08.010.

[76] Di G, Li Y, Zhu G, Guo X, Li H, Huang M, et al. Effects of acidification on the proteome during early development of Babylonia areolata. FEBS Open Bio 2019;9:1503–20. 10.1002/2211-5463.12695.

[77] Harms L, Frickenhaus S, Schiffer M, Mark F, Storch D, Held C, et al. Gene expression profiling in gills of the great spider crab Hyas araneus in response to ocean acidification and warming. BMC Genomics 2014;15:789. 10.1186/1471-2164-15-789.

[78] Guo Y, Zhou B, Sun T, Zhang Y, Jiang Y, Wang Y. An explanation based on energy-related changes for blue mussel Mytilus edulis coping with seawater acidification. Front Physiol 2021;12:761117. 10.3389/fphys.2021.761117.

[79] Gu M, Kong J, Di-Huang Peng T, Xie C, Yang K, et al. Molecular characterization and function of the Prohibitin2 gene in Litopenaeus vannamei responses to Vibrio alginolyticus. Dev Comp Immunol 2017;67:177–88. 10.1016/j.dci.2016.10.004.

[80] Choi K-M, Kim J-W, Kong HJ, Kim Y-O, Kim K-H, Park C-I. Molecular characterization, expression profiling, and functional analysis of prohibitin 1 in red seabream, Pagrus major. Fish Shellfish Immunol 2024;152:109770. 10.1016/j.fsi.2024.109770.

[81] Nelson WJ, Nusse R. Convergence of Wnt, β-catenin, and cadherin pathways. Science 2004;303:1483–7. 10.1126/science.1094291.

[82] Cadigan KM, Nusse R. Wnt signaling: a common theme in animal development. Genes Dev 1997;11:3286–305. 10.1101/gad.11.24.3286.

[83] Willert K, Brown JD, Danenberg E, Duncan AW, Weissman IL, Reya T, et al. Wnt proteins are lipid-modified and can act as stem cell growth factors. Nature 2003;423:448–52. 10.1038/nature01611.

[84] Hinck L, Nelson WJ. Wnt-1 modulates cell-cell adhesion in mammalian cells by stabilizing β-catenin binding to the cell adhesion protein cadherin. J Cell Biol 1994;124.

[85] Davis MA, Ireton RC, Reynolds AB. A core function for p120-catenin in cadherin turnover. J Cell Biol 2003;163:525–34. 10.1083/jcb.200307111.

[86] Drake JL, Schaller MF, Mass T, Godfrey L, Fu A, Sherrell RM, et al. Molecular and geochemical perspectives on the influence of CO2 on calcification in coral cell cultures. Limnol Oceanogr 2018;63:107–21. 10.1002/lno.10617.

[87] Wang X, Wang M, Wang W, Liu Z, Xu J, Jia Z, et al. Transcriptional changes of Pacific oyster Crassostrea gigas reveal essential role of calcium signal pathway in response to CO2-driven acidification. Sci Total Environ 2020;741:140177. 10.1016/j.scitotenv.2020.140177.

[88] Dineshram R, Xiao S, Ko GWK, Li J, Smrithi K, Thiyagarajan V, et al. Ocean acidification triggers cell signaling, suppress immune and calcification in the Pacific oyster larvae. Front Mar Sci 2021;8:782583. 10.3389/fmars.2021.782583.

[89] Wang T, Kong H, Shang Y, Dupont S, Peng J, Wang X, et al. Ocean acidification but not hypoxia alters the gonad performance in the thick shell mussel Mytilus coruscus. Mar Pollut Bull 2021;167:112282. 10.1016/j.marpolbul.2021.112282.

[90] Finkel T, Deng C-X, Mostoslavsky R. Recent progress in the biology and physiology of sirtuins. Nature 2009;460:587–91. 10.1038/nature08197.

[91] Vasquez MC, Tomanek L. Sirtuins as regulators of the cellular stress response and metabolism in marine ectotherms. Comp Biochem Physiol A Mol Integr Physiol 2019;236:110528. 10.1016/j.cbpa.2019.110528.

[92] Ding Q, Wang Y, Xia S-W, Zhao F, Zhong J-F, Wang H-L, et al. SIRT4 expression ameliorates the detrimental effect of heat stress via AMPK/mTOR signaling pathway in BMECs. Int J Mol Sci 2022;23:13307. 10.3390/ijms232113307.

[93] Vasquez MC, Beam M, Blackwell S, Zuzow MJ, Tomanek L. Sirtuins regulate proteomic responses near thermal tolerance limits in the blue mussels Mytilus galloprovincialis and Mytilus trossulus. J Exp Biol 2017:jeb.160325. 10.1242/jeb.160325.

[94] May MA, Tomanek L. Uncovering the roles of sirtuin activity and food availability during the onset of the heat shock response in the California mussel (Mytilus californianus): Implications for antioxidative stress responses. Comp Biochem Physiol B Biochem Mol Biol 2024;269:110902. 10.1016/j.cbpb.2023.110902.

[95] Chan SKN, Suresh S, Munday P, Ravasi T, Bernal MA, Schunter C. The alternative splicing landscape of a coral reef fish during a marine heatwave. Ecol Evol 2022;12:e8738. 10.1002/ece3.8738.

[96] Kędzierska H, Piekiełko-Witkowska A. Splicing factors of SR and hnRNP families as regulators of apoptosis in cancer. Cancer Lett 2017;396:53–65. 10.1016/j.canlet.2017.03.013.

[97] Tao Y, Zhang Q, Wang H, Yang X, Mu H. Alternative splicing and related RNA binding proteins in human health and disease. Signal Transduct Target Ther 2024;9:26. 10.1038/s41392-024-01734-2.

[98] Shi X, Zhang R, Liu Z, Zhao G, Guo J, Mao X, et al. Alternative splicing reveals acute stress response of Litopenaeus vannamei at high alkalinity. Mar Biotechnol 2024;26:103–15. 10.1007/s10126-023-10281-w.

[99] Shoshan Y, Liscovitch-Brauer N, Rosenthal JJC, Eisenberg E. Adaptive proteome diversification by nonsynonymous A-to-I RNA editing in coleoid cephalopods. Mol Biol Evol 2021;38:3775–88. 10.1093/molbev/msab154.

[100] Nishikura K. Functions and regulation of RNA editing by ADAR deaminases. Annu Rev Biochem 2010;79:321–49. 10.1146/annurev-biochem-060208-105251.

[101] Liscovitch-Brauer N, Alon S, Porath HT, Elstein B, Unger R, Ziv T, et al. Trade-off between transcriptome plasticity and genome evolution in cephalopods. Cell 2017;169:191-202.e11. 10.1016/j.cell.2017.03.025.

[102] Schulze-Osthoff K, Ferrari D, Riehemann K, Wesselborg S. Regulation of NF-κB activation by MAP Kinase cascades. Immunobiology 1997;198:35–49. 10.1016/S0171-2985(97)80025-3.

[103] Ninomiya-Tsuji J, Kishimoto K, Hiyama A, Inoue J, Cao Z, Matsumoto K. The kinase TAK1 can activate the NIK-IkB as well as the MAP kinase cascade in the IL-1 signalling pathway 1999;398.

[104] Hatada EN, Krappmann D, Scheidereit C. NF-κB and the innate immune response. Curr Opin Immunol 2000;12:52–8.

[105] Goodson MS, Kojadinovic M, Troll JV, Scheetz TE, Casavant TL, Soares MB, et al. Identifying components of the NF-κB pathway in the beneficial Euprymna scolopes-Vibrio fischeri light organ symbiosis. Appl Environ Microbiol 2005;71:6934–46. 10.1128/AEM.71.11.6934-6946.2005.

[106] Wang Y, Liu X, Wang W, Sun G, Feng Y, Xu X, et al. The investigation on stress mechanisms of Sepia esculenta larvae in the context of global warming and ocean acidification. Aquac Rep 2024;36.

[107] Kaplan DM, Largier JL, Navarrete S, Guiñez R, Castilla JC. Large diurnal temperature fluctuations in the nearshore water column. Estuar Coast Shelf Sci 2003;57:385–98. 10.1016/S0272-7714(02)00363-3.

[108] Smith KA, Rocheleau G, Merrifield MA, Jaramillo S, Pawlak G. Temperature variability caused by internal tides in the coral reef ecosystem of Hanauma bay, Hawai’i. Cont Shelf Res 2016;116:1–12. 10.1016/j.csr.2016.01.004.

[109] Fidopiastis PM, Von Boletzky S, Ruby EG. A new niche for Vibrio logei, the predominant light organ symbiont of squids in the genus Sepiola. J Bacteriol 1998;180:59–64.

