## Supplementary Figures for "Developmental and transcriptomic responses of Hawaiian bobtail squid early stages to ocean warming and acidification"

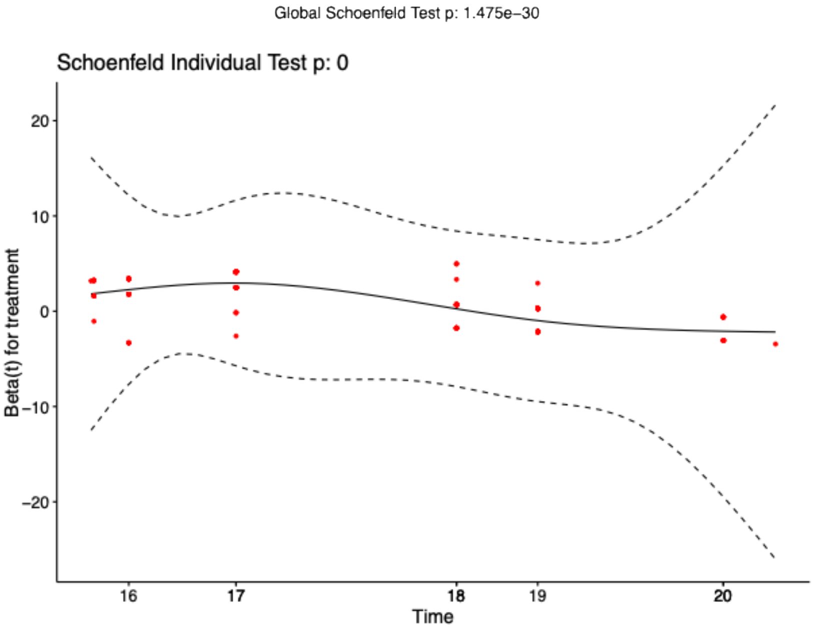


***Supplementary Figure 1*** – *Smoothed spline plots of Schoenfeld residuals of the hatching success Cox mixed effects model relative to time. Since the p-value is significant (p-value<0.05), the conditions of the Cox mixed effects model are not fulfilled. We followed the analysis of the data with a non-parametric model.*

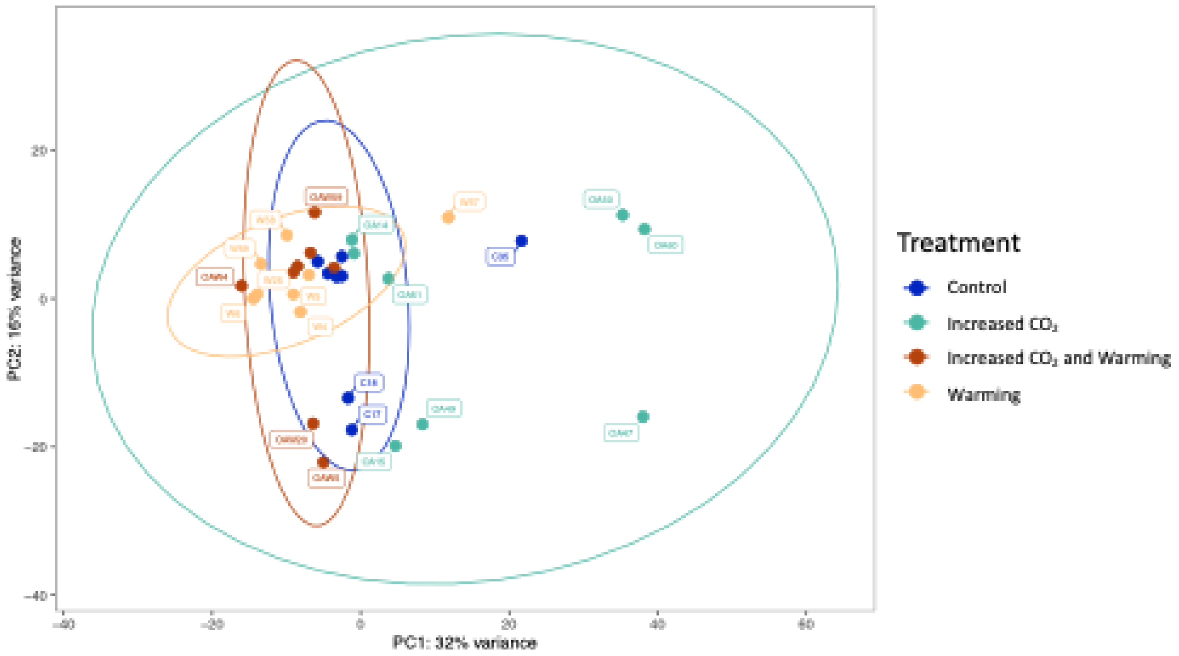


***Supplementary Figure 2*** – *Principal Component Analysis on the normalized count data of all samples for each treatment. The ellipses represent the 95% confidence interval. The dots are the data point (n=8 per treatment), the name of some samples are represented in the label (C='Control'; OA='High $CO_2$'; OAW='High $CO_2$ and warming'; W='Warming'). Outliers were identified by any data point outside the confidence ellipse: W57 and C35 were removed from the analysis.*

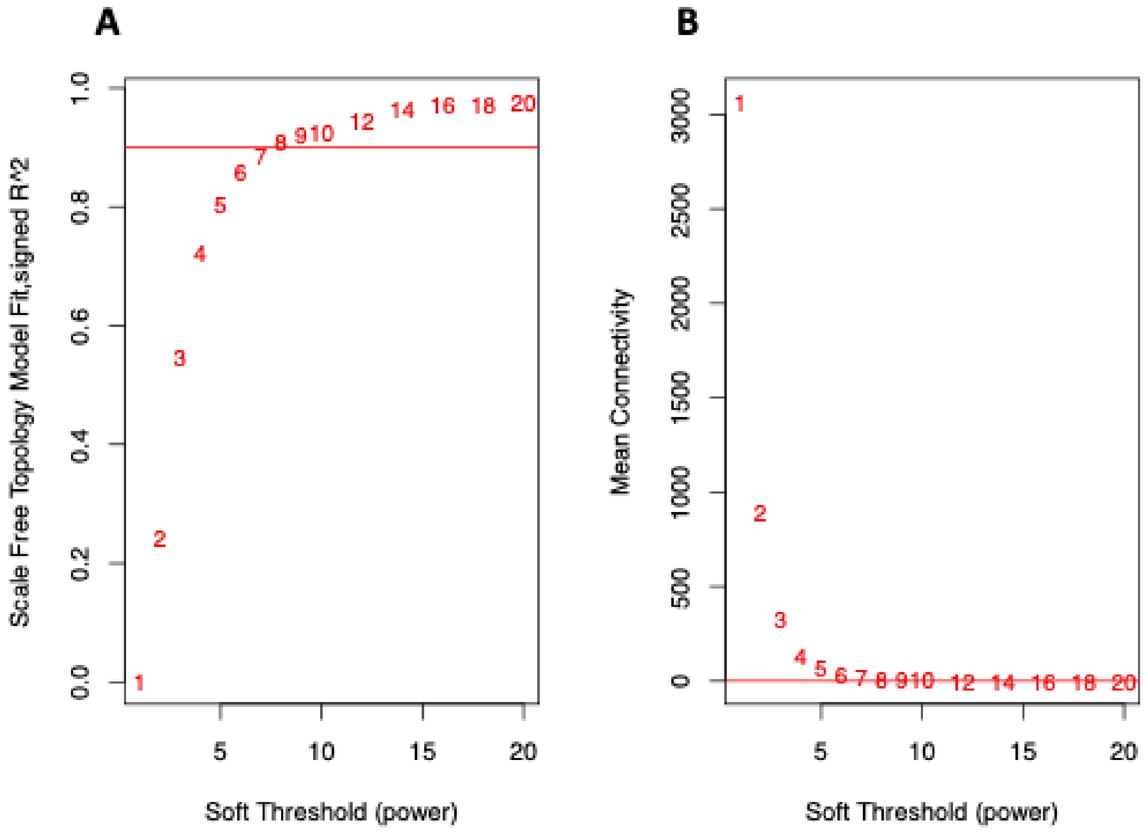


***Supplementary Figure 3*** *– Soft-threshold power for network construction during the Weighted gene co-expression network analysis (WGCNA). (A) Scale-free topology index in function of the soft-threshold. The network reaches a scale-free topology ($R^2$ > 0.9) when the soft-threshold is 8. Moreover, (B) the mean connectivity drops to 0 when the soft-threshold is 8.*

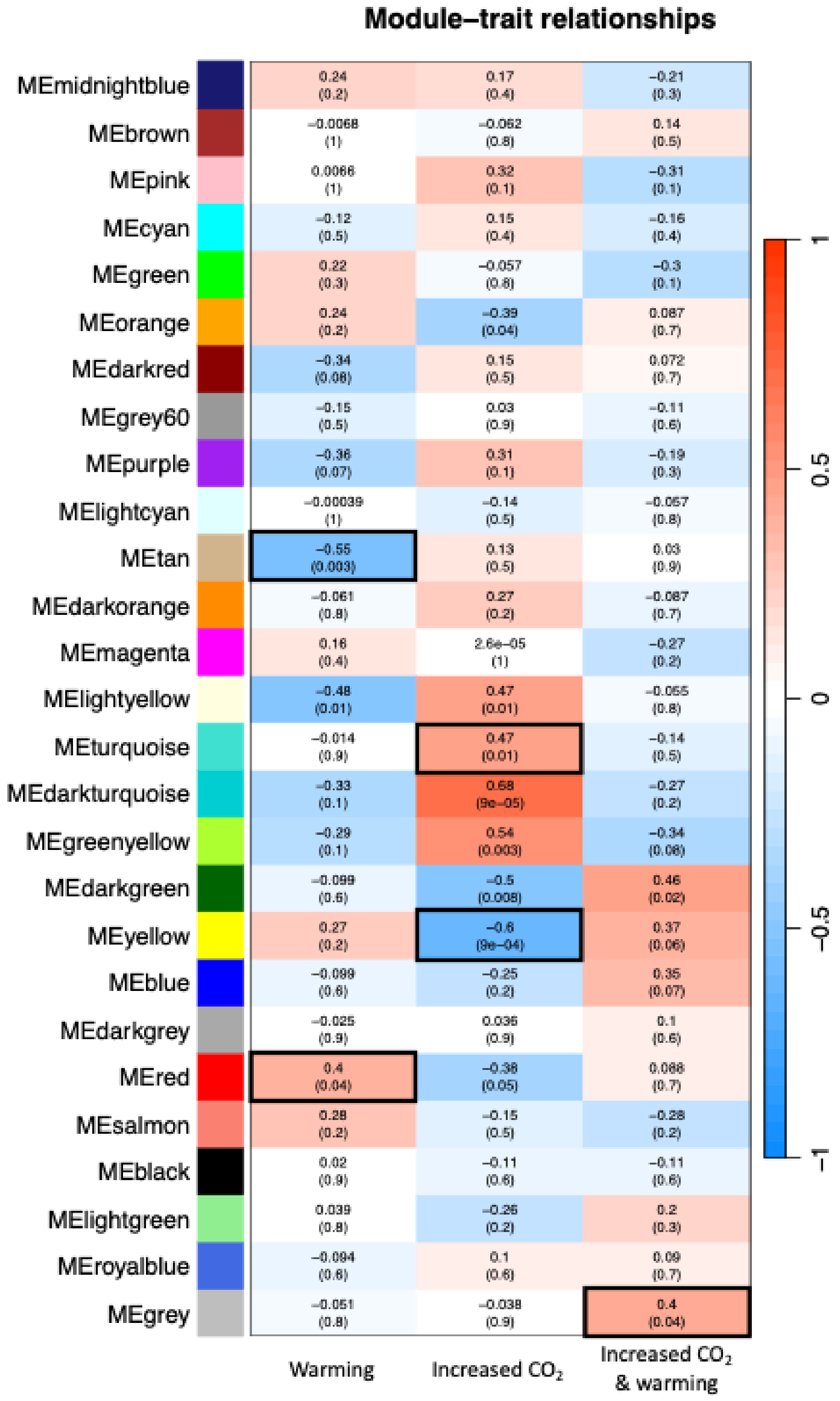


***Supplementary Figure 4*** *– Correlation heatmap between the modules eigengenes in function of the treatments, using the Pearson correlation test. Student asymptomatic p-value were calculated for the correlations and are represented under brackets, below the correlation value. Modules eigengenes were considered significantly correlated to a treatment when the p-value < 0.05. An increased expression of a module eigengenes, either positive or negative, within a given treatment reveals a relatively close connection between the module and the treatment. Red = positively correlated modules, while blue = negatively correlated modules.*
